# Cellular and Fibrillar Collagen Analyses in an Animal Model of Retinal Detachment-Related Proliferative Vitreoretinopathy Reveals a Defined Transition to Chronic Fibrosis

**DOI:** 10.1101/2022.12.14.520462

**Authors:** Cornelia Peterson, Clayton P. Santiago, Yuchen Lu, Antoinette Price, Minda M. McNally, William Schubert, Seth Blackshaw, Charles G. Eberhart, Mandeep S. Singh

## Abstract

**Purpose:** Proliferative vitreoretinopathy (PVR) is the most common cause of failure of surgically repaired rhegmatogenous retinal detachment (RRD). Chemically-induced and cell-injection PVR models do not fully simulate the clinical characteristics of PVR in the post-RRD context. There is an unmet need for translational models in which to study mechanisms and treatments specific to RRD-PVR.

**Methods:** RRD-PVR was induced in adult Dutch Belted rabbits. Posterior segments of enucleated globes were fixed or processed for RNA-Seq at 6 hours and 2, 7, 14, and 35 days post-induction. Histochemical staining and immunolabeling for glial fibrillary acidic protein (GFAP), alpha smooth muscle actin (αSMA), vascular endothelial growth factor receptor 2 (VEGFR2), CD68, and retinal pigment epithelium 65 kDa protein (RPE65) were performed, and labeling intensity was scored. Single cell RNA sequencing was performed.

**Results:** Acute histopathologic changes included intravitreal and intraretinal hemorrhage, leukocytic vitritis, chorioretinitis, and retinal rarefaction. Chronic lesions showed retinal atrophy, gliosis, fibrotic subretinal membranes, and epiretinal fibrovascular proliferation. Fibrillar collagen was present in the fibrocellular and fibrovascular membranes in chronic lesions. Moderate to strong labeling of glia and vasculature was detected in chronic lesions. At day 14, most cells profiled by single cell sequencing were identified as Müller glia and microglia, consistent with immunolabeling. Expression of several fibrillar collagen genes were upregulated in chronic lesions.

**Conclusions:** Histologic and transcriptional features of this rabbit model simulate important features of human RRD-PVR, including the transition to chronic intra and periretinal fibrosis. This high-fidelity *in vivo* model of RRD-PVR will enable further research on targeted treatment interventions.

---

Proliferative vitreoretinopathy (PVR), characterized by retinal contraction and periretinal proliferation, is the most common cause of failure after rhegmatogenous retinal detachment (RRD) repair. ^1–6^ There is currently no effective pharmacologic treatment or prophylaxis agent available for this surgical complication, and the cellular components and molecular drivers of PVR are incompletely elucidated. However, inflammation, the release of cytokines, altered expression of growth and transcription factors, and induction of epithelial-to-mesenchymal transition (EMT) of retinal pigmented epithelial (RPE) cells, fibroblasts, and glia have all been suggested as key factors. ^7–20^ Proposed targets for intervention include ligands, agonists, and inhibitors of cytokine and growth factor signaling pathways, but there remains an unmet need for effective PVR therapies and appropriate animal models of disease in which to evaluate them.^12,18,21–29^

The pathogenesis of RRD-PVR has historically been characterized by five developmental stages. The stages are: disruption of the blood-retinal barrier and resulting hypoxia, leukocyte chemotaxis and migration to the site of injury, EMT and proliferation of epiretinal, intraretinal, and subretinal cells, remodeling of the extracellular matrix (ECM) with membrane production, and fibrotic contraction. ^2–4,6,30,31^ However, the proportion and distribution of cell types involved and temporal and spatial regulation of these processes are poorly understood. The objective of the current study was to more comprehensively identify pertinent cell populations and describe the histologic lesions of PVR in a refined rabbit model of RRD-PVR. Here we present the cell types and ECM features of disease progression in RRD-PVR and demonstrate acute changes (primarily inflammation and hemorrhage) through the transition to epiretinal fibrovascular and subretinal fibrocellular membranes and chronic gliosis.

## Materials & Methods

### PVR Induction

This study was approved by the Johns Hopkins University’s Institutional Animal Care and Use Committee (protocol number: RB20M263). All animal experiments were performed in accordance with the guidelines for the Use of Animals in Ophthalmic and Vision Research of the Association for Research in Vision and Ophthalmology (ARVO) and in adherence with the ARRIVE 2.0 Guidelines.^32^ Adult female Dutch Belted rabbits (1.5-2.5kg) obtained from RSI (Robinson Services Inc, Mocksville, NC) were maintained in a temperature-controlled, 12h light cycle environments with *ad libitum* food and water. Briefly, animals were anesthetized with ketamine (35mg/kg IM) and xylazine (5mg/kg IM), intubated, and maintained on isoflurane. A refined approach to unilateral induction of RRD-PVR was achieved via lensectomy, pars plana vitrectomy, and retinotomy followed by retinal cryotherapy and intravitreal autologous platelet-rich plasma injection (*n* = 3 rabbits/time point); contralateral globes, receiving only enrofloxacin (5mg/kg subconjunctivally once postoperatively), lidocaine (1mg/kg subconjunctivally once postoperatively), and dexamethasone SP (0.5mg/kg subconjunctivally once postoperatively), served as controls.^33^

### Histology & Immunocytochemistry

Animals were humanely euthanized 6 hours and 2, 7, 14, and 35 days following RRD-PVR induction. Posterior segments of enucleated globes were harvested and then either fixed in 10% neutral-buffered formalin or 4% paraformaldehyde in PBS or stored at -80°C for later processing. Routine H&E staining of 5μm-thick FFPE and PFA-fixed, OCT-embedded sections were used to evaluate full-thickness retinal morphology and morphology of inner retina with associated fibrovascular membranes, respectively. Masson’s trichrome staining, performed on all FFPE sections by the JHMI Reference Histology Laboratory, was used to localize collagen expression. Immunohistochemical staining for retinal pigment epithelium 65 kDa protein (RPE65) was performed on all FFPE sections, and immunofluorescent staining for glial fibrillary acidic protein (GFAP), alpha smooth muscle actin (αSMA), vascular endothelial growth factor receptor 2 (VEGFR2), and CD68 was performed on all frozen sections using standard protocols and details as presented in **Supplementary Table 1**. Tissue sections were evaluated using an Olympus BX50 (Tokyo, Japan) and an Olympus AX70, respectively. The percentage of cells with positive labeling was estimated, and labeling intensity was evaluated to generate a semi-quantitative staining score (0: absent; 1: weak, focal; 2: moderate, multifocal; 3: strong, diffuse) independently by two pathologists (CP, CGE).

### RNA Sequencing

Extracted retinas (*n* = 2 per time point) were dissociated into single cells and processed to evaluate transcription profiling using single cell RNA-Seq (10X Genomics, Pleasanton, CA); UMAP analysis was performed to profile cell populations at day 14 post-induction and compared to controls as described elsewhere. ^34^ For heatmap analysis, collagen expression z-scores were obtained from log-transformed and pseudo-bulked transcript counts in control retina and lesions harvested at days 14 and 35, respectively.

### Statistical Analyses

The mode of semiquantitative immunolabeling scores was determined using Prism 8 GraphPad (v. 9.2.0; San Diego, CA) and recorded over time. The scRNA-Seq data was analyzed using R statistical language (R Core Team v. 4.1.0; Vienna, Austria). Default parameters in Seurat’s FindMarkers function was used to find differentially expressed genes using a non-parametric Wilcoxon rank sum test, and adjusted *p* values were obtained from Bonferroni corrections applied to all genes in the dataset (α = 0.05).

## Results

### Distinct histopathologic features of acute, subacute, and chronic lesions

We classified histopathologic changes as acute (6 hours – 2 days), subacute (7 days), and chronic (14 days and beyond). Retinas obtained from control animals demonstrated no significant hemorrhage, necrosis, or inflammation (**Fig. 1A-B**). Acute histopathologic changes (hour 6 and day 2) included intravitreal, intraretinal, and choroidal hemorrhage, infiltration of the vitreous, retina, choroid, and orbital soft tissues by large numbers of both degenerate and non-degenerate heterophils and fewer numbers of macrophages. Heterophils are leukocytes in rabbits which are functionally similar to neutrophils in humans.^35–38^ Rarefaction, edema, and fibrin deposition of the inner nuclear layers was also observed in the acute phase (**Fig. 1C-F)**. Sections of necrotic foci were present with swollen, hypereosinophilic ganglion cells containing pyknotic nuclei and disruption of photoreceptor outer segments in regions overlying subretinal fibrinosuppurative exudates.

**Figure 1:**
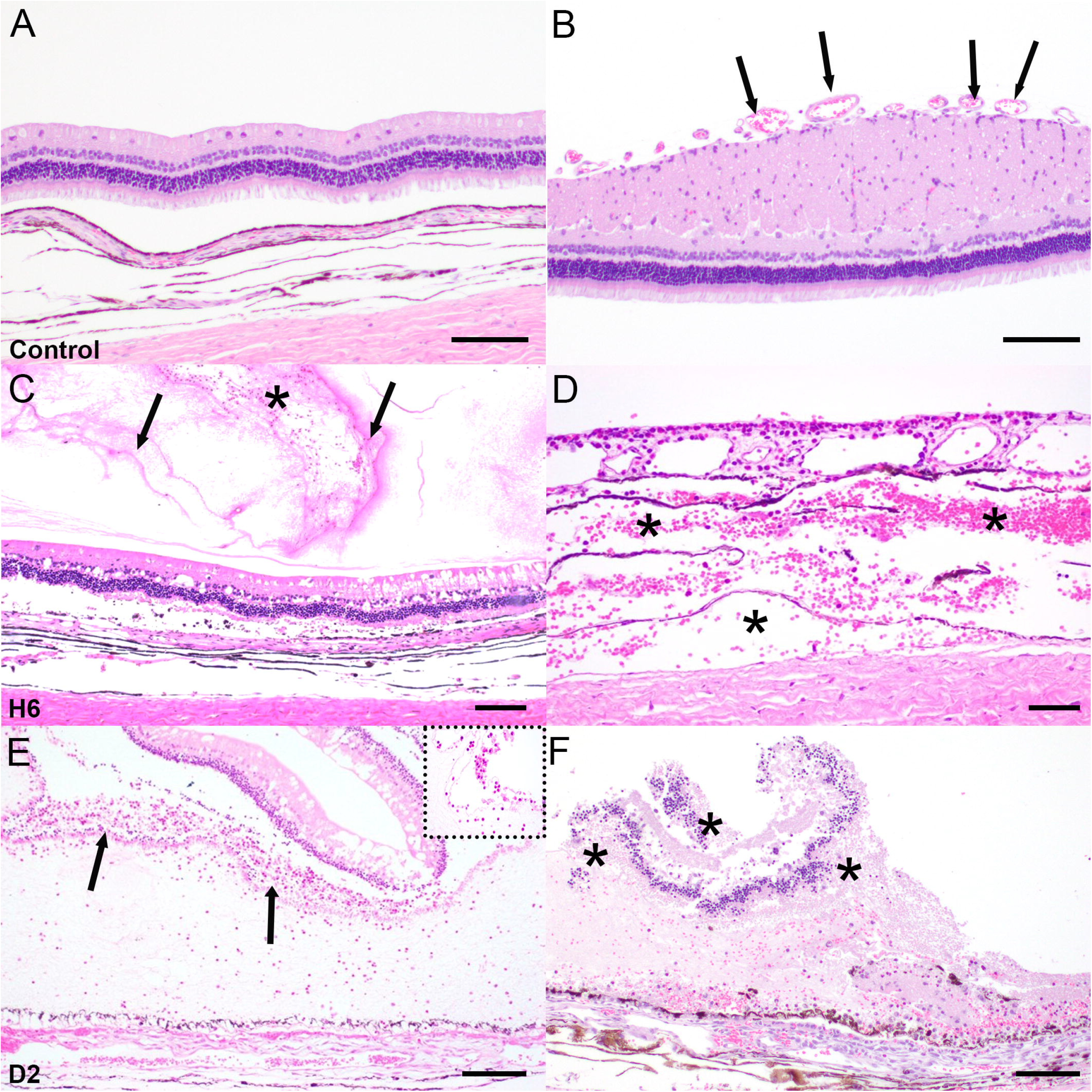
Representative histology of the normal control retina and acute histopathologic lesions following unilateral induction of RRD-PVR. No significant hemorrhage, inflammation, or rarefaction were observed within the control neural retina (A) or the medullary streak (B; arrows indicate surface vasculature). Moderate to severe vitreal hemorrhage (asterick) and fibrin exudation (arrows) (C) and moderate to severe choroidal hemorrhage (asterices) (D) were observed at 6 hours post-induction. Acute heterophilic inflammation within the subretinal space (arrows) (E) with inflammatory cells shown at higher magnification (inset) and foci of necrosis of the neural retina (asterices) (F) were observed 2 days post-induction. Scale bar: 50μM.

Subacute changes observed at day 7 included full-thickness retinal folds and persistent edema and rarefaction of the inner retina admixed with infiltrating microglia. Vitreal hemorrhage had nearly resolved at this time point; however, a mixed population of inflammatory cells remained in the vitreous and subretinal space, and erythrophagocytosis was observed. Early atrophic change, particularly of the outer retina, and gliosis were also present by day 7 (**Fig. 2A-B**). Chronic lesions (days 14 and 35) included panretinal atrophy and progressive gliosis, highlighted by prominent radial Müller glia (**Supplementary Fig. 1**). Subretinal fibrocellular membranes, often with anterior dispersion of single RPE cells through the inner retina, and epiretinal fibrovascular membranes in the region of the medullary streak were also observed at later time points (**Fig. 2C-F**).

**Figure 2:**
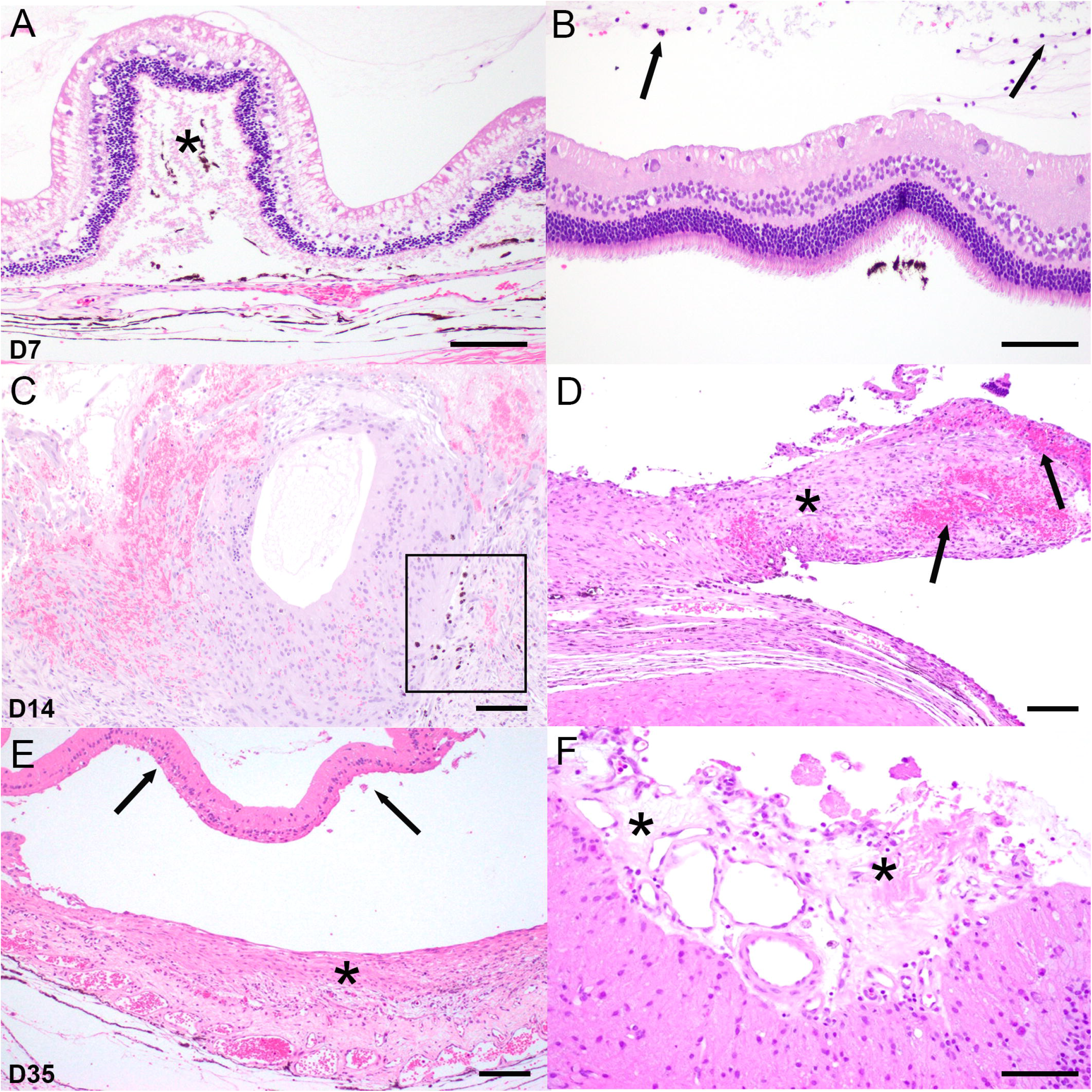
Representative histology of subacute and chronic histopathologic lesions following unilateral induction of RRD-PVR. Progressive rarefaction of the inner neural retinal layers along with full-thickness retinal folds (asterick) (A) and moderate infiltration of the vitreous by a mixed population of inflammatory cells (arrows) (B) were present at day 7 post-induction. Resolving hemorrhage, moderate to severe atrophy, and full-thickness folding of the neural retina with anterior migration of pigmented RPE cells (boxed area) (C) and the formation of subretinal fibrocellular membranes (asterick) with intramembranous hemorrhage (arrows) (D) were observed at day 14 post-induction. At day 35 post-induction progressive, dense fibrous subretinal membranes (asterick) were attached to the RPE and Bruch’s membrane posterior to the detached and atrophic retina (arrows) (E). Dense epiretinal fibrovascular membranes (asterices) were also observed in the region of the medullary streak (F). Scale bar: 50μM.

### Progressive accumulation of fibrillar collagen in epiretinal and subretinal membranes

To assess the location and extent of epi- and subretinal membranes containing a collagenous component, we conducted trichrome staining. Retinas with acute and subacute lesions were trichrome negative; however, trichrome-positive strands of fibrillar collagen were observed within the vitreous at day 2 and day 7. Semiquantitative analysis showed that moderate trichrome positivity began at day 14 and progressed by day 35 (**Fig. 3A-G**). Chronic lesions exhibited trichrome-positive fibrillar collagen in the subretinal fibrocellular and epiretinal fibrovascular membranes.

**Figure 3:**
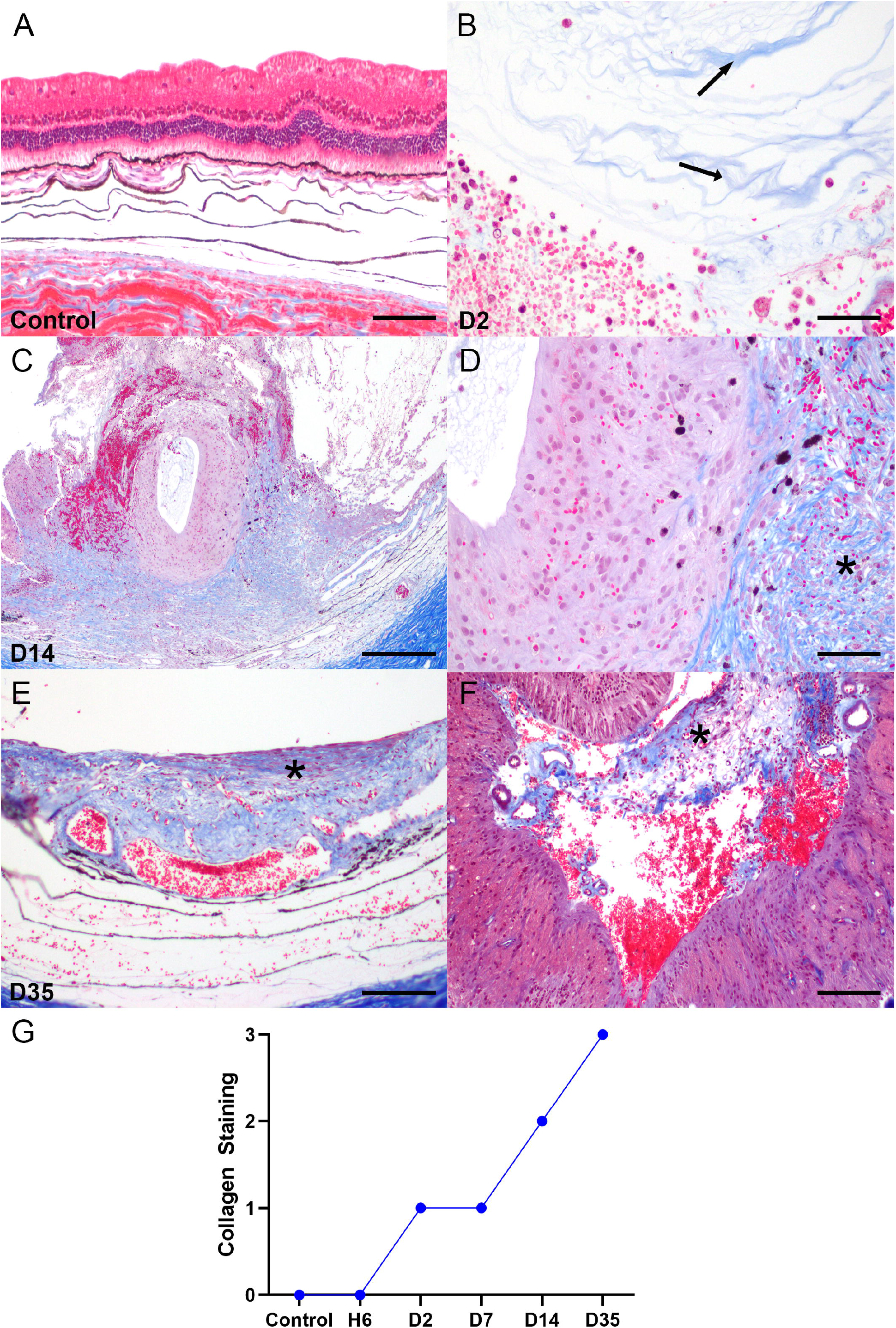
Expression of fibrillar collagen in the rabbit retina following unilateral induction of RRD-PVR. No collagen was observed in the vitreous, retina, or subretinal space in the control tissue (A: Masson’s trichrome stain, Scale bar: 50μM). Thin strands of fibrillar collagen (blue) were observed in the vitreous in association with acute heterophilic vitritis by day 2 (arrows) (B, Scale bar: 25μM). By day 14, the subretinal space was expanded by fibrillar collagen that abutted the folded and gliotic neural retina (C, Scale bar: 100μM). Higher magnification demonstrating intraretinal gliosis and subretinal fibrosis (asterick) (D, Scale bar: 50μM). By day 35, the subretinal membranes (asterick) contained increasing amounts of fibrillar collagen with fewer cellular components (E, Scale bar: 100μM), and the epiretinal fibrovascular membranes of the medullary streak (asterick) comprised a dense collagenous matrix (F, Scale bar: 50μM). The intensity of trichrome staining increased over time (G; *n* = 3 per time point).

We hypothesized that multiple collagen species were involved in peri-retinal membrane formation. To interrogate gene expression of diverse collagen species, we used single cell RNA sequencing. Heatmap analysis for fibrillar collagen revealed that *COL1A1* was upregulated in Müller glia at day 14 post-injury relative to control samples, while *COL5A1* and *COL5A2* were upregulated at day 35 post-induction relative to controls (**Fig. 4A**). These data suggest that multiple collagen species may be involved in the formation of pathological PVR membranes in promoting of peri-retinal membrane contractility.

**Figure 4:**
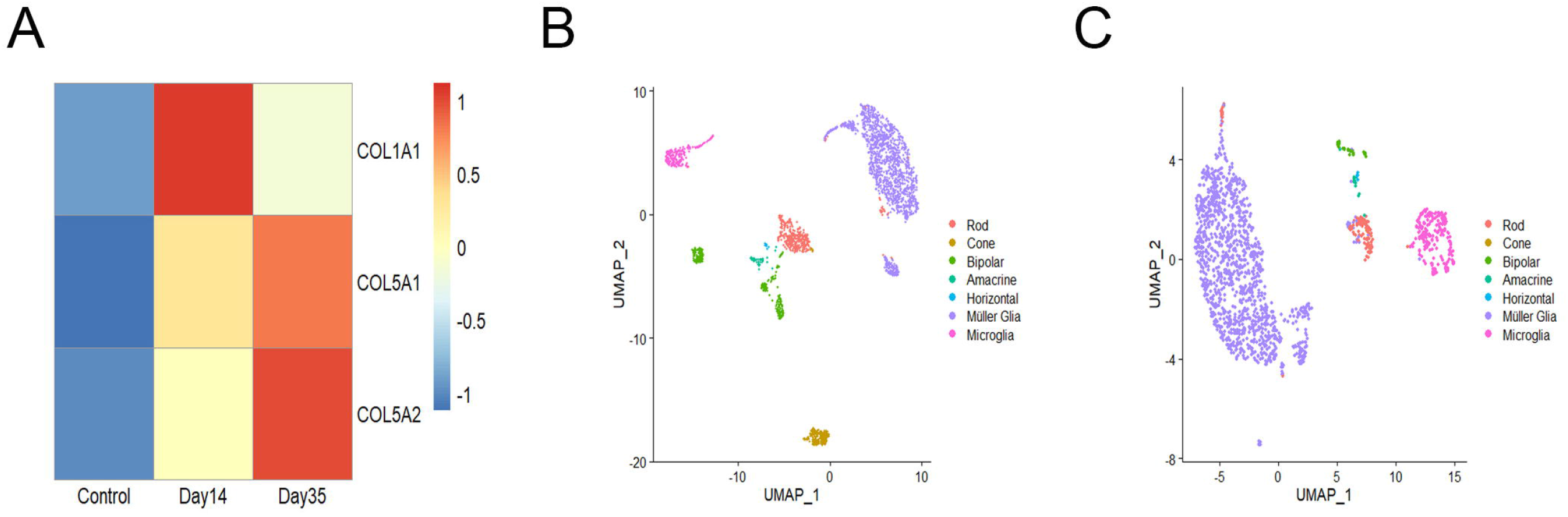
scRNA-Seq of the rabbit retina following unilateral induction of RRD-PVR. Heatmap for differentially-expressed fibrillar collagen produced by Müller glia demonstrating upregulation in chronic lesions (A; *n* = 2 per time point). UMAP plots for single cell RNA-Seq of control rabbit retinas (B) and day 14 post unilateral induction of RRD-PVR (C). Note the change in the number of cells (Müller glia and microglia) over time.

### Temporal expression of VEGF Receptor 2 in fibrovascular lesions

We observed epiretinal fibrovascular proliferation in the region of the medullary streak in chronic RRD-PVR. To assess the extent and temporal pattern by which VEGF signaling may play a role in cell proliferation in the context of these lesions, we stained acute, subacute, and chronic specimens for VEGFR2. In acute and subacute specimens, VEGFR2 was weakly expressed. However, expression of VEGFR2 associated with epiretinal fibrovascular membranes in the medullary streak peaked at day 14 (**Fig. 5A-D**). There was only moderate expression at day 35.

**Figure 5:**
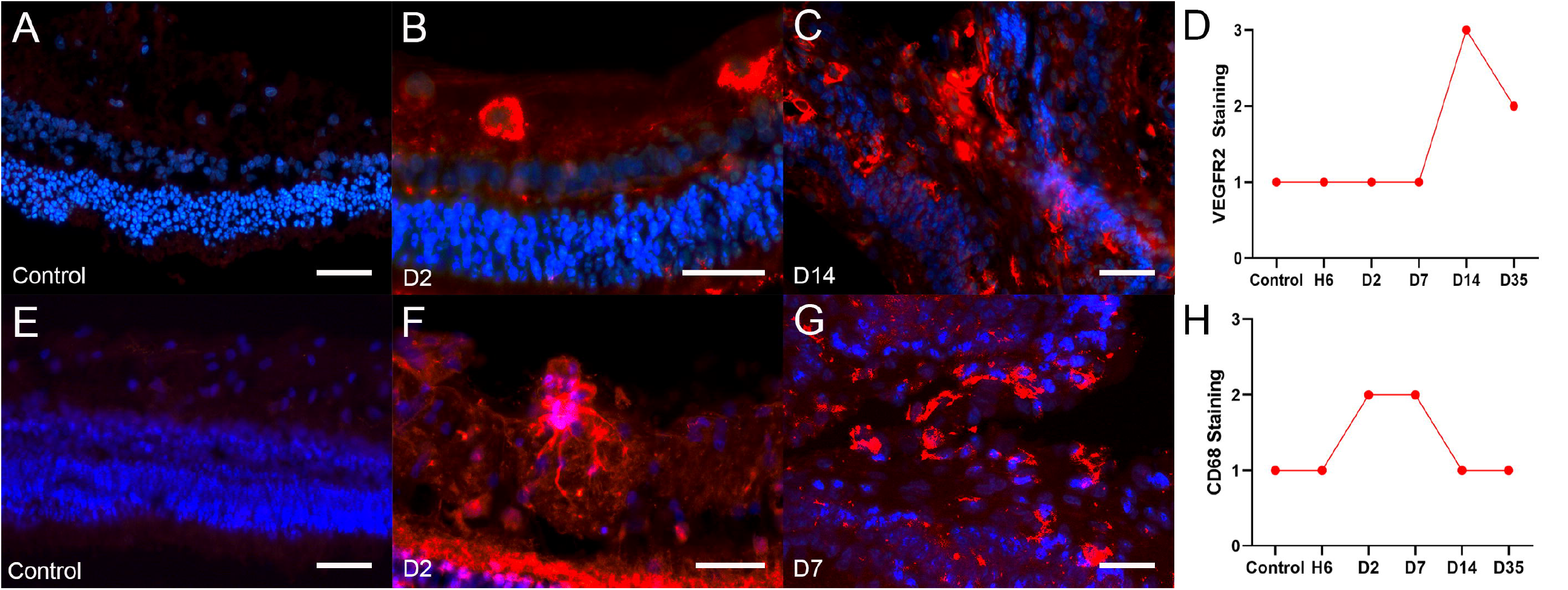
Expression of VEGFR2 and CD68 in the rabbit retina following unilateral induction of RRD-PVR. No labeling of vessels was observed within the neural retina (A: Red: VEGFR2, Blue: DAPI, Scale bar: 25μM) in the control tissue or at day 2 or day 7 in any region other than the medullary rays (B, Scale bar: 10μM). By day 14 there was a marked increase in the number or small vessels within the inner layers of the medullary streak and in association with epiretinal fibrovascular membranes (C, Scale bar: 25μM). VEGFR2 staining was increased at day 14 and but began to decrease at day 35 (D; *n* = 4 per time point). Weak labeling of microglia was observed within the neural retina in the control tissue (E: Red: CD68, Blue: DAPI, Scale bar: 25μM). By day 2, there was moderate expression of CD68 by cells morphologically consistent with microglia within the inner layers of the neural retina (F, Scale bar: 25μM). At day 14 there was moderate expression of CD68 by microglia associated with epiretinal fibrovascular membranes in the medullary streak (G, Scale bar: 25μM). CD68 staining peaked at day 2 and day 7 (H; *n* = 3 per time point).

### Microglia and gliotic changes

To assess the abundance of microglial cells, we combined CD68 immunolabeling and morphologic features, with a small proportion of cells having abundant cytoplasm and a larger eccentric nucleus, a morphology supportive of circulating macrophages; however, the majority of these cells were smaller with rarefied processes, consistent with microglia. In acute and subacute lesions, CD68 expression, associated with increased numbers of retinal microglia, peaked at days 2 and 7. CD68+ cells were located primarily in the inner retinal layers and epiretinal membranes of the medullary streak (**Fig. 5E-H**). At chronic time points, CD68+ cells were observed at an intensity and proportion similar to controls. We assayed upregulated expression of GFAP to evaluate Müller glia hypertrophy. GFAP+ Müller glia were restricted to the neural retina (**Fig. 6B-D**). At day 14 post-induction, relatively few neurons were captured by UMAP analysis, and the majority of cells profiled were identified as Müller glia and microglia (**Fig. 4B-C**).

**Figure 6:**
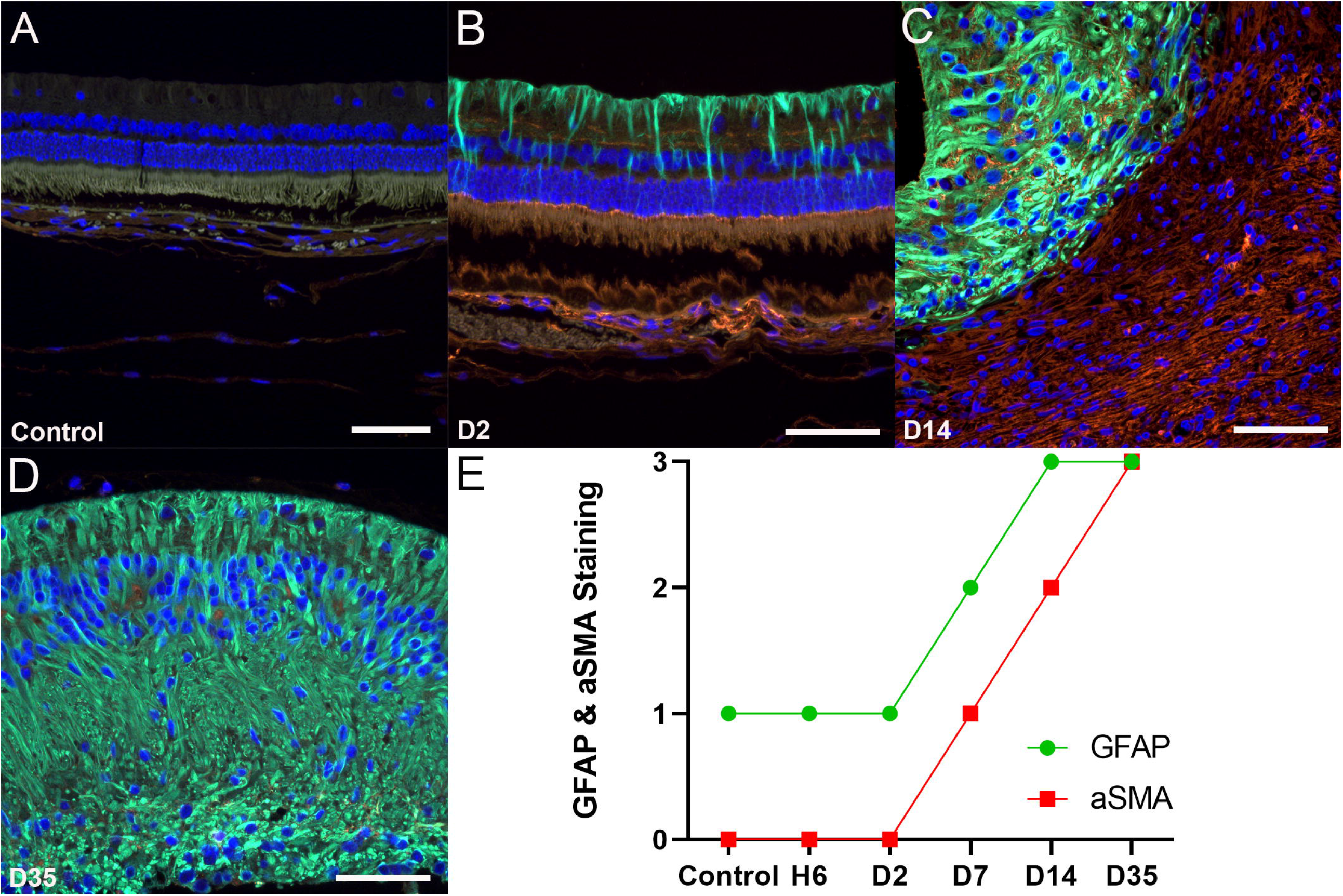
Expression of GFAP and αSMA in the rabbit retina following unilateral induction of RRD-PVR. Weak expression of GFAP by Müller glia within the neural retina was observed in the control globes. αSMA was not expressed in controls (A: Red: αSMA, Green: GFAP, Blue: DAPI, Scale bar: 50μM). By day 2, increased GFAP expression by Müller glia was observed within the neural retina (B, Scale bar: 50μM). At day 14, GFAP was strongly expressed by proliferating and hypertrophied Müller glia within the neural retina, and αSMA was strongly expressed within subretinal fibrocellular membranes (C, Scale bar: 50μM). At day 35, gliosis and GFAP expression were diffuse wtihin the neural retina (D, Scale bar: 50μM). The staining of both GFAP and αSMA increased over time (E; *n* = 4 per time point).

### Subretinal RPE de-differentiation and proliferation

We used RPE65 as a marker of differentiated RPE cells. RPE65-positive cells were attenuated (weak to absent) at day 14 relative to controls, with expression loss corresponding to regions of subretinal fibrosis and foci of pigmented cells observed migrating anteriorly through the subretinal membranes toward the interface with the gliotic neural retina (**Fig. 7A-D**). At day 14, upregulated expression of αSMA within the subretinal fibrocellular membrane indicated dedifferentiation and EMT of RPE cells (**Fig. 6C**).

**Figure 7:**
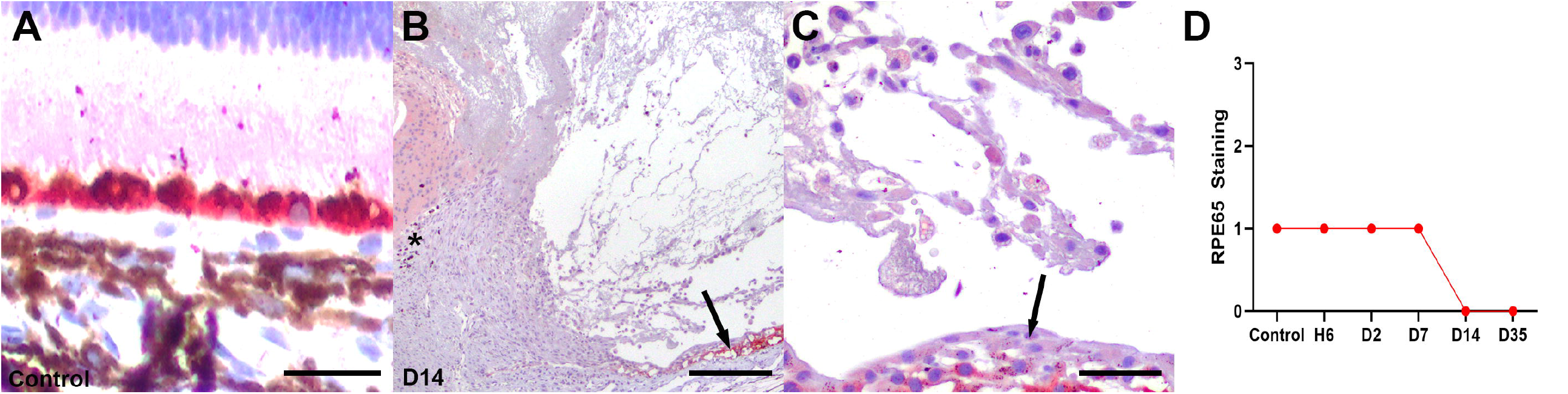
Expression of RPE65 in the rabbit retina following unilateral induction of RRD-PVR. RPE65 expression was observed in the RPE in the control retinas with normal cuboidal architecture (A: Red: RPE65, Scale bar: 25μM). At day 14, RPE65 expression was largely absent in the region of the subretinal membrane (arrow) and in foci of migrating pigmented cells (asterick), consistent with dedifferentiation of RPE (B, Scale bar: 100μM). Higher magnification highlighting the loss of RPE65 expression in RPE that have undergone EMT and transitioned from normal cuboidal epithelial cells and assumed a flattened, fibroblastic appearance (arrow). Moderate numbers of subretinal heterophils and macrophages are present, several of which contain red bloods cells within the cytoplasm (erythrophagocytosis) (C, Scale bar: 25μM). RPE65 staining decreased over time (D; *n* = 4 per time point).

## Discussion

These data indicate that the rabbit model of rhegmatogenous PVR is broadly analogous to human PVR. The epi- and subretinal fibrocellular membranes in this model appear to simulate similar lesions that are commonly found in clinical PVR. While the rabbit is a commonly utilized animal model for PVR due to the large size of the globe and its ability to manifest features similar to the human condition, existing models have not consistently demonstrated both epi- and subretinal membranes, the key histologic features of human PVR.^16,29,39–46^ In addition, our data demonstrate the involvement of multiple collagen species in the development of PVR membranes, suggesting that multiple profibrotic signaling pathways may be involved. These rabbit data also validate the presence of additional features of human PVR, i.e., increased microglial invasion into epiretinal membranes, Müller glia activation, and RPE de-differentiation.^47^

This manuscript also addresses two main knowledge gaps in the rabbit PVR models, namely acute changes (6 hours and 2 days) and epi-versus subretinal fibrosis. We focused on acute timepoint analysis because the early pathological changes represent potential treatment targets to prevent more severe retinal damage. Regarding subretinal fibrosis, we felt this key feature has received inadequate attention despite being an important – and surgically challenging, in terms of curative intervention – component in the most severe cases of human PVR.

Earlier studies have primarily focused on epiretinal membrane formation. Chen *et al* reported disruption of the inner limiting membrane and epiretinal fibrosis in their model after 10 days post lesion induction. ^16^ In their dataset, critical acute changes occurring earlier than 10 days could not be identified, and data on subretinal membrane formation was not presented.^16^ Similarly, histologic sections presented in Hirose *et al* and Khoroshilova-Maslova *et al* demonstrated tractional retinal folds with epiretinal membranes at four- and eight-weeks post-induction.^39,41^ Epiretinal membranes and RPE proliferation were reported at 30 days post-induction but not at more acute time points, and subretinal membranes were also not targeted for analysis.^39,41^

Other investigators report variable degrees of retinal thickening, gliosis, and atrophy. ^29^ Elevated αSMA expression in epiretinal membranes at time points ranging from 22 days to 11 weeks post-induction have been reported, but subretinal data are unavailable. ^29,42–46^ Khanum *et al* reported evaluation of their rabbit model one and 30 days post-induction; however, it is unclear from their figures from which time points their photomicrographs of H&E-stained or immunolabeled sections were obtained.^40^ Our 6-hour and 2-day data broadly align with the findings of Zahn *et al* at days 3 and 7 post-induction, in which they assayed Müller glia, microglia, and macrophages.^48^

Early inflammatory changes, characterized by leukocyte infiltration and local cytokine and growth factor production, have long been considered the nidus for vascularized epiretinal membrane formation in PVR, with mononuclear cells and histiocytes more often implicated in traumatic etiologies. ^49–52^ In the current study, we demonstrate fibrinohemorrhagic and necrotizing vitritis and chorioretinitis with subsequent heterophilic infiltration by 6 hours and day 2 post-induction, respectively. Increasing expression of CD68 within the inner retinal layers of the medullary streak was observed in the subacute period, peaking at days 2 and 7 post-induction, with a lag before the development of neovascularization in the fibrovascular epiretinal membranes observed at day 14. Collectively, these results suggest that early damage or ischemia is characterized initially by acute infiltration by heterophils, the subsequent release of proinflammatory cytokines and growth factors, which then, in turn, promote microglial recruitment and neovascularization. Under hypoxic conditions, reduced ubiquination or TNFα-mediated induction of HIF-1α promotes transcriptional activation of hypoxia responsive elements, leading to increased expression of VEGFR2 to restore tissue perfusion.^53,54^ The contribution of microglia to the retinal inflammatory response and regulation of retinal and choroidal vasculature has been well-documented, further supporting a role in neovascularization for the microglia observed in association with the fibrovascular epiretinal membranes in the current study.^19,55^

Jin *et al* described a key transition in the course of human PVR at day 30 when highly cellular, loosened ECM becomes more densely fibrotic.^31^ In the current study, we validate this transition in ECM remodeling of both the epiretinal and subretinal membranes at day 35 post-induction with compaction and paucicellularity of the membranes when compared to day 14. Our scRNA-Seq data confirmed this differential expression of fibrillar collagens over time with upregulation of *COL1A1, COL5A1*, and *COL5A2*, at days 14 and 35 post-induction, respectively. Further, this transition corresponded to an increase in αSMA and fibrillar collagen expression in both the epi- and subretinal membranes as demonstrated with immunolabeling and Masson’s trichrome staining, respectively. The transition also correlated with increased Müller glia proliferation and hypertrophy as demonstrated by GFAP immunolabeling when compared to controls. GFAP and vimentin are commonly utilized as markers of retinal gliosis in the context of PVR, and Eastlake *et al* reported that numerous inflammatory mediators are expressed by Müller glia *in vitro* and upregulated in the gliotic retina.^17,18,56^

In the current study, we demonstrate loss of RPE65 expression by RPE cells by day 14 post-induction, corresponding to the same time point that subretinal fibrocellular membrane formation was earliest observed. Additionally, rare pigmented cells, that were negative for RPE65 labeling and therefore likely to correspond to dedifferentiated RPE cells or possibly choroidal melanocytes, were observed at the anterior margin of these subretinal membranes, abutting the gliotic neural retina. Together, these findings suggest dedifferentiation of RPE cells in response to chronic injury before transforming to a more mesenchymal cell type, and support the notion the migration of proliferative RPE cells not only contributes to epiretinal membranes, but also subretinal fibrocellular membranes, in PVR.^57^

Failure to adequately sample RPE and subretinal tissue for dissociation and RNA isolation, precluded this critical cell population from subsequent sequencing analyses and the opportunity to confirm the dedifferentiation during EMT, observed histologically at day 14 post-induction, and represents a limitation of the current study. A further limitation is that the layer-specific distribution (i.e., epi-versus sub-retinal membrane) of collagen expression is unclear from the current dataset because retinal specimens, including all membranes, were processed together for scRNA-seq.

## Conclusions

In summary, we have demonstrated the cellular and ECM features throughout the time course of PVR development using a novel rabbit model of RRD-PVR. Additionally, using RNA-Seq, we have confirmed the relevant cell populations present at a critical transition presented at day 14 post-induction and identified fibrillar collagen species that are differentially expressed by Müller glia. High-fidelity *in vivo* models for RRD-PVR, such as that presented here, are essential for future investigations of targeted treatment interventions.

## Acknowledgments

This project was funded by Bayer AG as part of a sponsored research collaboration with the Wilmer Eye Institute, Johns Hopkins Hospital. Dr. Peterson’s and Dr. Santiago’s salary support was provided by NIH T32 OD011089 (PI: Mankowski) and NIH 2T32EY007143 (PI: Zack), respectively.

---

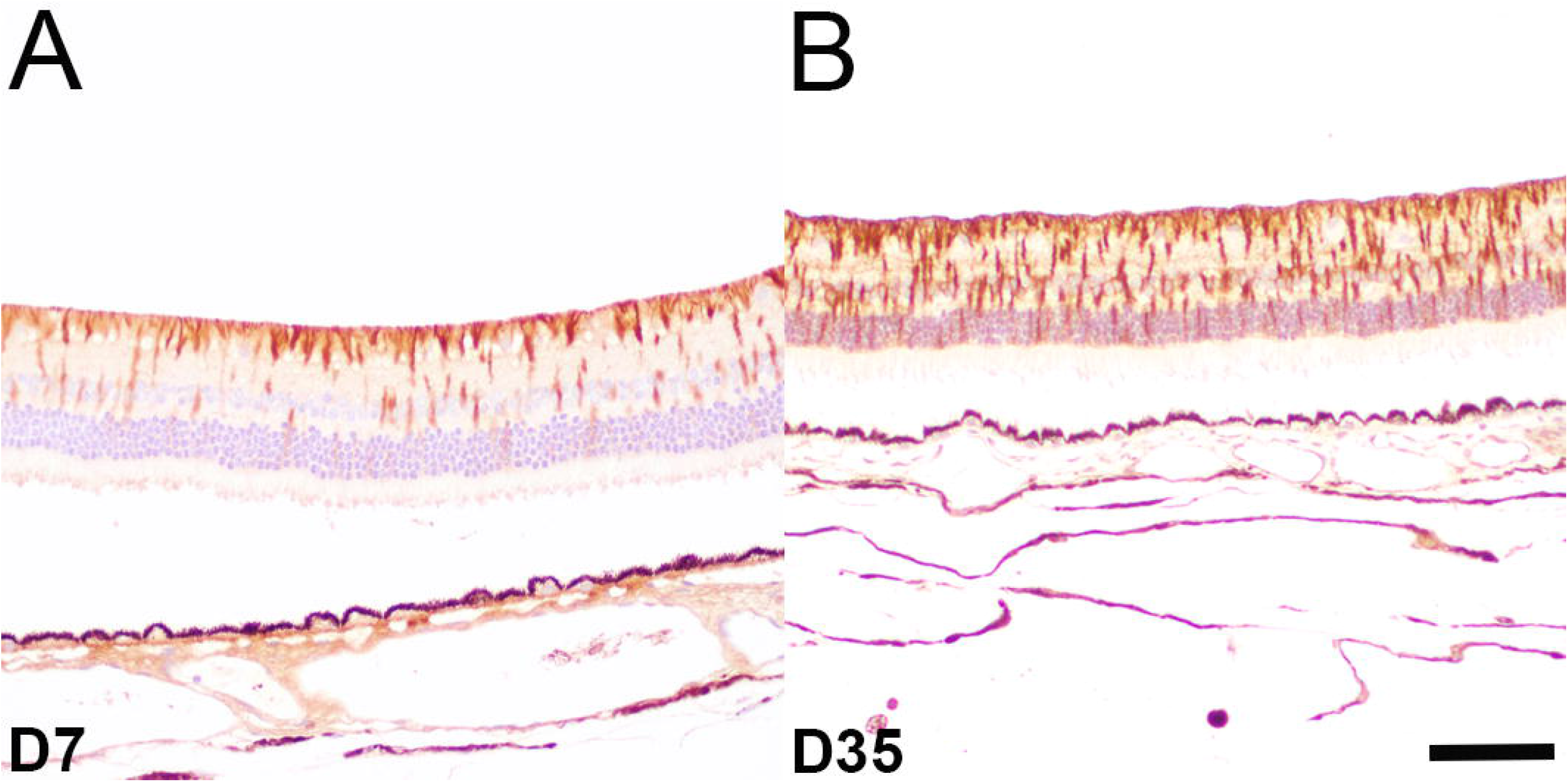

