## Supplemental Table 1 for "Cellular and Fibrillar Collagen Analyses in an Animal Model of Retinal Detachment-Related Proliferative Vitreoretinopathy Reveals a Defined Transition to Chronic Fibrosis"

**Supplementary Materials**

**Supplementary Table 1: Reagents used for immunolabeling.**

| Antibody/Reagent | Company (Location) | Catalog # | Concentration |
|---|---|---|---|
| AEC Chromogen | Abcam (Cambridge, UK) | ab103742 | NA |
| AlexaFluor 488 Anti-mouse IgG (H+L) | Cell Signaling Technology (Danvers, MA) | 4408 | 1:500 |
| AlexaFluor 555 Anti-rabbit IgG (H+L) | Cell Signaling Technology | 4413 | 1:500 |
| αSMA | Invitrogen Life Technologies (Waltham, MA) | 710487 | 1:5000 |
| CD68 | Cell Signaling Technology | 76437 | 1:100 |
| DAB Chromogen | Thermo Fisher Scientific (Waltham, MA) | ICN980681 | NA |
| GFAP (IF) | Invitrogen Life Technologies | 14-9892-82 | 1:250 |
| GFAP (IHC) | Agilent (Santa Clara, CA) | Z033429-2 | 1:200 |
| RPE65 | Invitrogen Life Technologies | MA1-16578 | 1:500 |
| Vectastain Elite Mouse IgG ABC-HRP Kit | Vector Laboratories (Burlingame, CA) | PK-6102 | NA |
| Vectastain Elite Rabbit IgG ABC-HRP Kit | Vector Laboratories | PK-6101 | NA |
| VEGFR2 | Cell Signaling Technology | 2479 | 1:200 |

**Supplementary Figure 1: Expression of GFAP in the rabbit retina following unilateral induction of RRD-PVR.** Moderate upregulation of GFAP (brown) was observed in the foot processes of actived Müller glia along the inner limiting membrane at day 7 post-induction (A), and panretinal staining was further increased at day 35 post-induction (B). Scale bar: 50μM.
